# Benchmarking Cell-Type-Specific Spatially Variable Gene Detection Methods Using a Realistic and Decomposable Simulation Framework

**DOI:** 10.1101/2025.11.26.690782

**Authors:** Weiqi Li, Xinzhou Ge, Yuan Jiang

**Affiliations:** Department of Statistics, Oregon State University, Corvallis, OR, USA, 97331

**Keywords:** Benchmarking, Spatial transcriptomics, Simulation, Spatially variable genes, Visium, Xenium

## Abstract

Identifying spatially variable genes within individual cell types is essential for characterizing spatially organized cell states and microenvironments from spatial transcriptomics data. Several computational methods have been developed for identifying cell-type-specific spatially variable genes (ctSVGs), but their relative performance and practical utility under realistic biological complexity remain largely unknown. To address this gap, we present the first systematic benchmark study of all five existing ctSVG detection methods—CELINA, STANCE, C-SIDE, CTSV and spVC—using an integrated evaluation framework that combines idealized simulations, Xenium-based realistic simulations, and a decomposition-based diagnostic analysis. We compared the methods in terms of detection accuracy, scalability, and usability. Across realistic datasets generated on various tissue types, all methods experienced sharp declines in detection accuracy and substantial inflation of false discoveries compared to idealized simulations. To explain this failure, we developed a new simulation framework that decomposes the “realness” of the realistic simulation into interpretable biological and technical components, enabling us to attribute method-specific performance losses to specific components, including realistic diversity of cell types, heterogeneous cell layouts, null gene distributions, capture efficiency and realistic intra-cell-type spatial patterns. Together, our results show that no single method dominates across detection accuracy, scalability and usability, and we further clarify why current ctSVG methods fall short in realistic settings. We summarize these tradeoffs into a practical user guide to support method selection and highlight key challenges in developing robust, scalable ctSVG detection tools for real spatial transcriptomics data.

## 1. Background

The spatial transcriptomics (ST) technologies have advanced rapidly in recent years, enabling gene expression profiling within intact tissues while preserving the spatial information at various resolutions, from spot-level measurements encompassing dozens of cells [1] to near-cellular resolutions [2] and even subcellular measurements [3]. The spatial dimension added to transcriptomic data enables unprecedented insights into complex tissue organization and biological mechanisms such as tumor microenvironments [4], developmental trajectories [5], and cell-to-cell interactions [6]. One common task for all ST profiles is to identify spatially variable genes (SVGs) that display non-random spatial patterns across the tissues [7–9]. SVGs capture biologically informative signals and are critical to downstream analysis including dimension reduction, delineating spatial domains, and characterizing tissue organization [10, 11]. Numerous computational methods have been developed for detecting SVGs, and they have also been extensively reviewed and benchmarked in prior studies [12–16]. One common challenge in SVG detection is that cell types are unevenly distributed across the tissue, making it difficult to distinguish true spatial heterogeneity within cell types from spatial variation driven by differences in expression levels across cell types. As a result, the detected SVGs can be dominated by cell-type marker genes, and their spatial patterns are largely affected by cell-type composition patterns. This limitation motivates the study of cell-type-specific spatially variable genes (ctSVGs)—genes that exhibit non-random spatial variation within specific cell types. ctSVGs extend the concept of SVGs into finer resolution and enable deeper biological interpretation, revealing spatially organized cell states, signaling programs, disease-associated microenvironments, and subpopulations within cell types.

To date, five computational methods have been proposed to detect ctSVGs, and all of them model gene expression using regression frameworks that incorporate both cell-type constant effects and spatial effects. These methods fall into two categories: (1) fixed-effect models including C-SIDE [17], CTSV [18], and spVC [19], which represent spatial variation through fixed coefficients on constructed spatial kernels or spatial basis functions, and (2) random-effect models including CELINA [20] and STANCE [21], which model spatial variation through kernel-based covariance matrices. With multiple methods developed within the past three years and more expected to come, it has become increasingly challenging for users to determine which approach to use for identifying ctSVGs.

Although existing reviews of SVG detection methods have included some of the ctSVG detection methods, these reviews only provide high-level summaries and lack quantitative bench-marking [12, 13]. While individual method papers include comparisons with other alternatives, these comparisons are limited in scope, involving only limited methods and evaluation criteria. More importantly, existing methods have used unrealistic simulation datasets to compare their performance with others, typically containing only a small number of cell types, uniform cell-type composition, homogeneous gene expression distribution, and predefined spatial patterns. Such *idealized simulations* fail to capture the biological complexity observed in real tissues, and therefore the comparison results obtained from these simulations cannot accurately reflect the performance of different methods on real data. Recent benchmarking studies for SVG detection methods have used reference-based simulators such as scDesign3 [22] and SRTsim [23] to generate more realistic simulated datasets [14–16]. However, no existing work has included *realistic simulations* specifically for benchmarking ctSVG detection methods, which involve additional challenges. One key challenge in defining ground truth ctSVGs is to distinguish true spatial patterns within an individual cell type from spatial variations driven by differences in cell-type compositions.

Usability and scalability are also important considerations when selecting ctSVG detection methods, and users often need guidance in these areas. Although individual methodological papers have raised concerns about limitations in usability or computational efficiency, there has been no systematic summary or comprehensive guideline to help practitioners compare different methods along these dimensions. Collectively, these gaps highlight an emerging need for a comprehensive benchmarking study based on *realistic simulations*, and a practical user guide that clarifies method strengths, limitations, and usability in realistic settings.

In this work, we benchmarked all five existing computational methods for identifying ctSVGs: C-SIDE, CTSV, spVC, CELINA, and STANCE. To investigate their detection performance beyond simplified settings, we developed a simulation pipeline that generates Visium-like spot-resolution datasets with realistic cell-type composition and ground truth ctSVGs derived from real-world ST data using scDesign3. Importantly, we also evaluated and confirmed the biological validity of this pipeline, ensuring that the simulated data preserve key spatial and transcriptional features observed in real tissues. We used multiple metrics to comprehensively evaluate method performance in terms of detection accuracy and statistical calibration. In *idealized simulations*, most methods achieved high detection accuracy but showed inadequate statistical calibration and poor false discovery control. In *realistic simulations*, we observed that detection accuracy declined substantially for all methods, accompanied by further inflation of false discoveries.

One key innovation of this study is the development of a new simulation framework, *decomposed simulation*, which decomposes the “realness” of the *realistic simulations* into interpretable components. These “realness” components include realistic tissue structure, such as numbers and proportions of cell types, and realistic gene expression, such as gene distributions and intra-cell-type spatial patterns. By generating datasets with various levels of “realness”, we performed controlled experiments to quantify the effect of each “realness” component on each method’s performance. The method-specific effects revealed by the *decomposed simulation* help explain the discrepancies between the results from the *idealized* and *realistic simulations* and provide detailed guidance for selecting methods suited to specific tissue contexts. In addition, this realistic and decomposable simulation framework offers a practical tool for future method developers to identify which aspects of realistic complexity most strongly affect method performance and to clarify the potential underlying causes of underperformance, offering insights into how to improve methods’ effectiveness in analyzing real spatial transcriptomic data.

## 2. Results

### 2.1. Comprehensive simulation framework for method benchmarking

A central challenge in benchmarking ctSVG detection methods is the lack of realistic datasets with ground truth. To address this challenge, we developed a unified benchmarking framework that systematically evaluates five ctSVG detection methods in terms of detection accuracy, statistical calibration, and computational scalability. Existing methods such as CELINA and STANCE [20, 21] have proposed relatively simple simulation designs: few cell types with balanced proportions, uniform cell density, homogeneous gene distributions, and strong, consistent spatial signals. We based our first set of simulations on these designs and refer to them as *idealized simulations*. As illustrated in Fig. 1b, these *idealized simulation* settings include diverse cell-type compositions and intra-cell-type spatial patterns, providing a useful foundation for evaluation of different methods. In Results 2.2, we present the simulation results from *idealized simulations*.

**Fig. 1:**
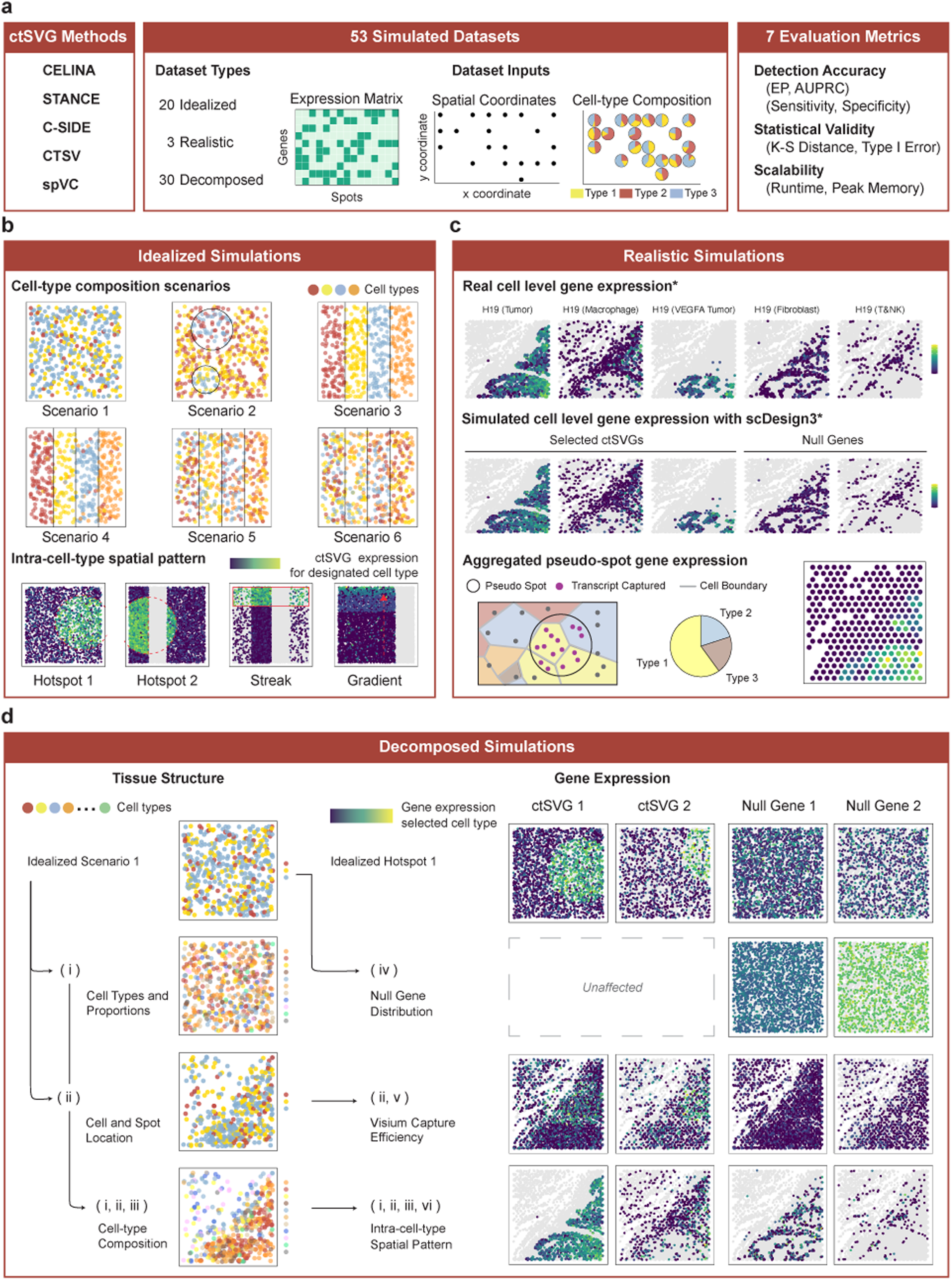
Overview of the benchmarking study for ctSVG detection methods. **a** Left: five benchmarked ctSVG detection methods. Middle: summary of the 53 simulation datasets across idealized, realistic, and decomposed settings, and illustration of the spot-level ST datasets. Right: evaluation metrics we used to benchmark the methods, covering detection accuracy, statistical calibration, and computational scalability. **b** Visualization of the *idealized simulations*, where the number of cell types is small, cell-type compositions are from the six predefined scenarios, and the spatial patterns of ground-truth ctSVGs are from the four predefined patterns. **c** Visualization of (Top) real-world ovarian cancer data and (Middle) simulated data in the realistic simulation. (Bottom) Workflow of generating *realistic simulation*, from real-world Xenium cell-level expression to scDesign3-simulated ctSVGs and null genes, followed by aggregation into Visium-like pseudo-spots. **d** Schematic of the *decomposed simulations*, showing the individual components of biological and technical complexity, how they are introduced to isolate their impacts on method detection performance, and visualization of the corresponding decomposed simulation dataset for each component or component combination.

However, these settings rely entirely on idealized spatial structures and fail to capture the complexity of real biological systems. To address these limitations, we used the scDesign3 framework which leverages real data as a reference to generate more realistic and biologically grounded simulations. We adapted this framework and developed a simulation pipeline that generates realistic Visium-like, spot-resolution datasets from subcellular-resolution measurements [22]. Specifically, we first modeled gene expression using subcellular-resolution data, which naturally provides reliable cell-type annotations and clear intra-cell-type spatial patterns. We then simulated gene expression at the cell level based on fitted gene expression models and aggregated cells into pseudo-spots while accounting for Visium capture efficiency (Fig. 1c). We refer to these simulation settings as *realistic simulations* and present the simulation results in Results 2.3.

In *realistic simulations*, the performance of all methods worsened notably compared with *idealized simulations*, making it difficult to draw meaningful conclusions in terms of detection accuracy and statistical calibration based on these poor performances. To understand the discrepancy between *idealized simulation* and *realistic simulation* results and to identify the factors driving the failure of each individual method in *realistic simulations*, we decomposed the “realness” of scDesign3-based simulations into two major aspects—tissue structure and gene expression—each containing three interpretable components (Fig. 1d). We then simulated datasets with varying levels of realness and compared method performances across these datasets. These controlled experiments diagnose which aspects of realness most strongly influence the detection performance of each method. We refer to these simulations as *decomposed simulations* and provide a detailed discussion on their results in Results 2.4. Finally, we examined computational scalability by varying the numbers of genes and spots and present the results in Results 2.5.

### 2.2. Detection accuracy and statistical calibration in *idealized simulations*

In *idealized simulations*, we simulated ctSVGs based on predefined intra-cell-type spatial patterns, e.g., hotspot, streak, or gradient (Fig. 1b). These simulations are idealized. Each dataset was first simulated at the single-cell level and aggregated into spots arranged on equal-sized square grids, assuming gene expression was unaffected by capture efficiency. Cell locations were drawn from a Poisson point process, resulting in uniform cell density across the tissue. Each dataset included three or four cell types with either balanced or mildly uneven proportions, and the cell-type composition was configured as either spatially uniform or varying across predefined domains (Fig. 1b). Gene counts were sampled from negative binomial distributions. Null genes had constant means across cell types and spatial locations. Marker genes exhibited fixed fold-change differences in their designated cell type while remaining at the null level in all others. ctSVGs also exhibited fixed fold-change differences in their designated cell type but only within predefined spatial domains while matching the background null level outside those domains.

#### 2.2.1. Most methods have high detection accuracy but are sensitive to cell-type proportion imbalance

Detection accuracy was evaluated using the area under the precision–recall curve (AUPRC) and early precision (EP). AUPRC is widely used in settings with rare positives but is sensitive to the prevalence of true positives. In contrast, EP measures the precision among the top *K* ranked detections, where *K* is the number of true ctSVGs, and is therefore invariant to prevalence of positives. To make detection performance more comparable across simulation settings with different signal prevalences, we primarily interpret performance differences using EP.

As shown in Fig. 2a, CTSV, CELINA, STANCE and C-SIDE all achieved high detection accuracy under balanced cell-type proportions. However, EP declined noticeably when cell-type proportions were mildly unbalanced (Fig. 2b). Under balanced cell-type proportions, CTSV and CELINA showed the strongest detection accuracy with an average EP of 95% and 91%, respectively. In some scenarios, CELINA nearly perfectly ranked ctSVGs over non-ctSVGs, but its quality-control filtering excluded around 8% of ground truth ctSVGs, placing a hard cap on attainable accuracy. Sharing a similar filtering mechanism, C-SIDE faced the same limitation. Results in Fig. 2b showed that STANCE had reduced EP for streak and gradient intra-cell-type spatial patterns because it relies on a single distance-based kernel — in contrast to CELINA’s eleven kernels, CTSV’s ten, and the spline bases used by C-SIDE and spVC. Notably, spVC’s detection accuracy is unstable even under balanced settings. When cell types were mildly unbalanced (e.g., 10%, 30%, 60% in Scenario 1 of Fig. 1b), detection accuracy of all methods dropped to around 50%. Under such unbalanced settings, all methods maintained strong EP above 66% for the dominant cell type comprising 60% of total cells, but performed poorly on the minority cell type comprising 10% of total cells, where EP ranged only from 7% to 40%.

**Fig. 2:**
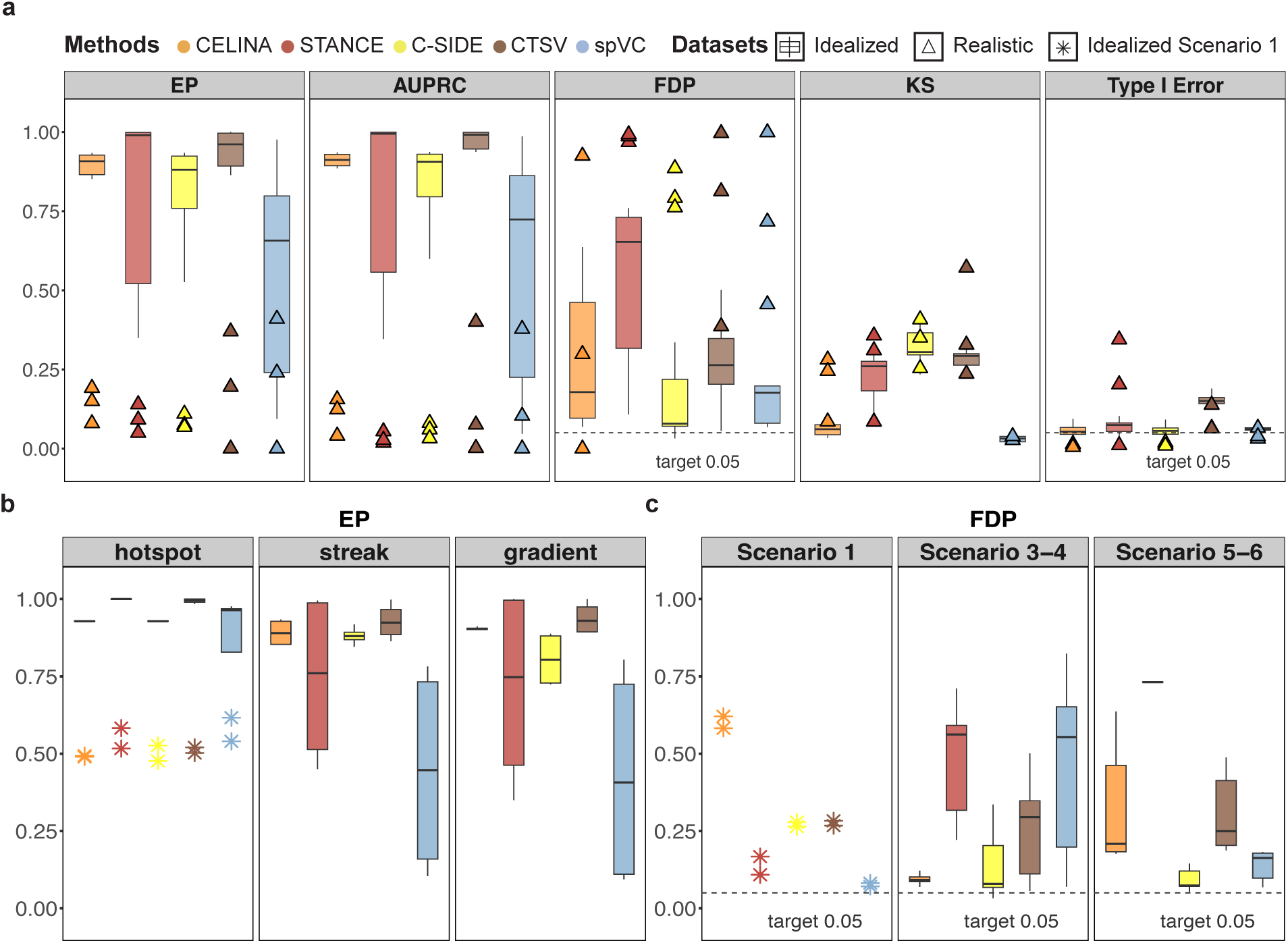
Benchmarking detection accuracy and statistical calibration using *idealized simulations* and *realistic simulations*. **a** Performance of five ctSVG detection methods across idealized datasets generated from six predefined scenarios (boxplots) and realistic datasets generated from three real tissue types (triangles) characterized by detection accuracy metrics (EP, AUPRC), error control (FDP), and statistical calibration (K-S distance, Type I error). **b** Boxplots of EP in *idealized simulations* stratified by intra-cell-type spatial patterns. The simulated datasets with Scenario 1 and Hotspot pattern are highlighted. **c** FDP in *idealized simulations* across three groups of composition scenarios.

#### 2.2.2. Most methods have inadequate statistical calibration, and none effectively controlled false discoveries even under idealized settings

We first evaluated the statistical calibration of the methods using Type I Error rate for non-ctSVGs at the nominal level of 0.05, as well as Kolmogorov-Smirnov (K-S) distance between the empirical *p*-value distribution for non-ctSVGs and the uniform distribution. Among all the five methods, CELINA and spVC achieved the best statistical calibration (Fig. 2a). All the other methods—including C-SIDE, CTSV, and STANCE—exhibited substantial deviations from the expected uniform distribution.

For both C-SIDE and CTSV, miscalibration largely stems from how they handle multiple tests for spatial effect of a single gene. Both methods conduct multiple tests across different kernels or basis functions for a single gene and then combine the resulting *p*-values. Specifically, C-SIDE computes as many *p*-values as the user-specified number of spline basis functions, applies Bonferroni correction across them, and takes the minimum adjusted *p*-value as the final output for the single spatial effect. This approach produces an average of 30% *p*-values exactly equal to 1 and thus far deviates from expected calibration (Supplementary Fig. S4). CTSV separately computes Cauchy-combined *p*-values for two sets of kernels, one based on the x-coordinate and the other based on the y-coordinate. It then reports the smaller of the two *p*-values, resulting in anti-conservative calibration.

STANCE is designed as a two-stage test, where only the genes found to have spatial pattern in the entire tissue in its stage 1 test proceed to the stage 2 ctSVG test. For non-ctSVGs tested in stage 2, STANCE produces anti-conservative *p*-values and thus displays a notable deviation from uniform distribution. STANCE still has anti-conservative calibration, with an average type I error of 8% among all non-ctSVGs including genes correctly excluded before conducting stage 2 test. The inflation was primarily driven by the model’s omission of cross-cell-type constant effects, where gene expression displays no intra-cell-type spatial patterns but differs by constant fold changes across cell types. Without accounting for these effects, STANCE mis-attributes spatial variation arising from non-uniform cell-type composition to true ctSVG signal, resulting in an average 68% type I error for cell-type marker genes that are non-ctSVG (Supplementary Fig. S3).

We then used false discovery proportion (FDP) at a target false discovery rate (FDR) of 5% to evaluate different methods’ abilities to control false discoveries. Despite CELINA’s well-calibrated *p*-values, it did not effectively control false discoveries, resulting in an FDP of 27%. The inflation was most pronounced when cell types overlap spatially as in the *idealized simulation* Scenarios 1, 5, and 6 rather than forming distinct domains as in Scenario 3 (Fig. 2c). In the extreme case of Scenario 1, where all three cell types uniformly scatter across the tissue, CELINA had an average FDP of 60%. Even under moderately overlapping conditions (Scenarios 5 and 6), its average FDP remained inflated at 33%.

### 2.3. Reduced detection accuracy and inflated false discoveries in *realistic simulations*

We generated three realistic datasets based on real Xenium subcellular-resolution spatial data from three distinct tissue types and disease states: breast tumor, ovarian cancer and healthy lymph node (Fig. 3a). For each real dataset, we first simulated cell-level count data using scDesign3, fitting the gene expression model separately for each cell type to preserve realistic expression variation within cell types. To emulate the Visium capture process, we then randomly sampled about 50% of the transcripts and aggregated them into pseudo-spots (Fig. 1c). The proportion of transcripts sampled was calibrated using the proportion of gene transcripts that fall within the Visium capture discs for each cell in the real data.

**Fig. 3:**
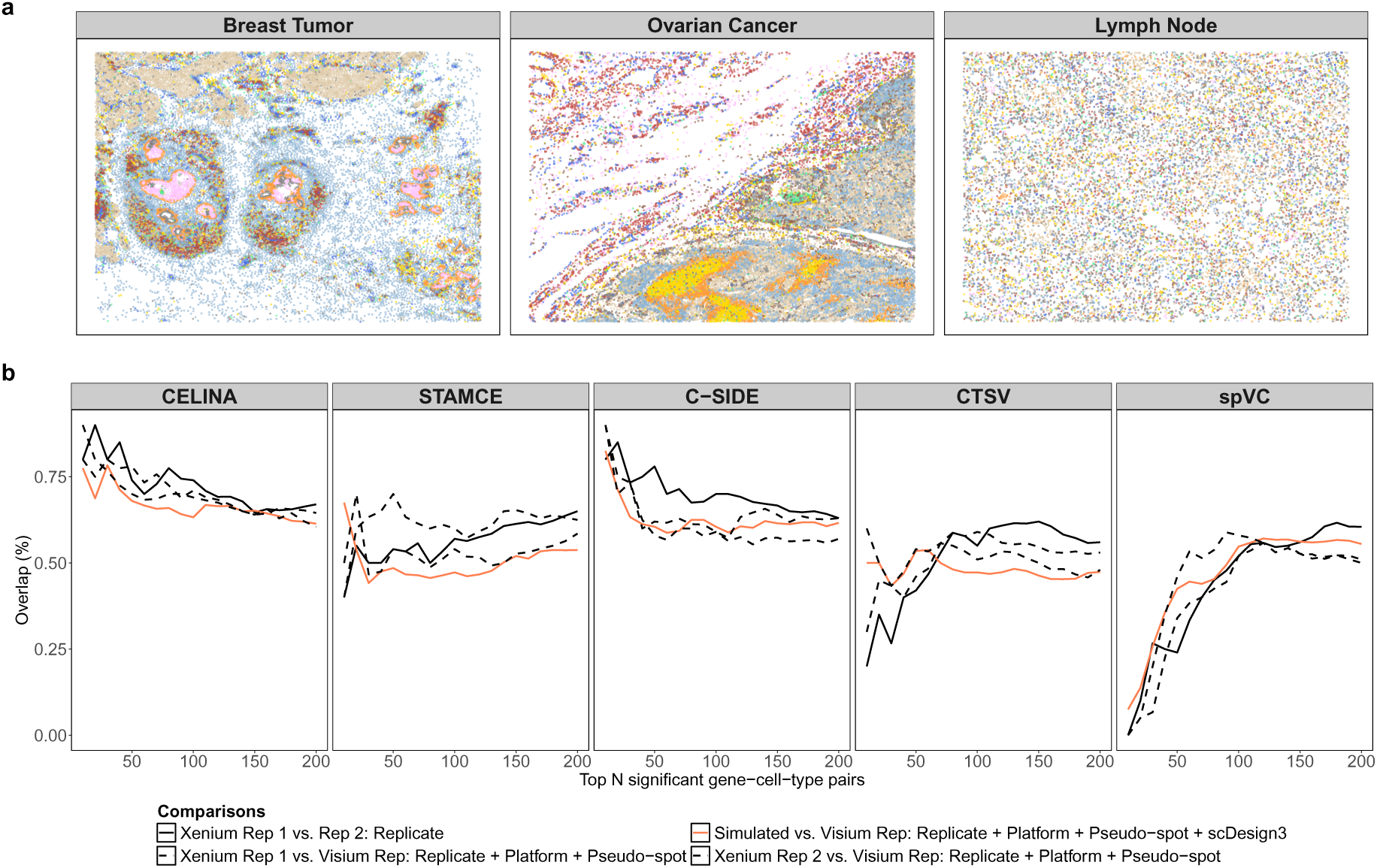
Real tissue structures and validation of the biological realism of the realistic simulation. **a** Visualization of the spatial organization of three tissues used for generating *realistic simulations*, with colors denoting cell types. Breast tumor and ovarian cancer display heterogeneous cell density and spatially varying cell-type composition, whereas lymph node tissue exhibits more uniform density with scattered voids. **b** Agreement of detected ctSVGs across datasets. The y-axis is the proportion of overlapping ctSVGs between the discoveries from two datasets in each comparison, and the x-axis is the number of top significant discoveries. For each method, the solid line represents the comparison between pseudo-spot datasets derived from two Xenium serial replicates; the orange line represents the comparison between the simulated datasets and the real Visium dataset from a matched serial section; the two dashed lines represent comparisons between each of the pseudo-spot datasets derived from two Xenium replicates and the real Visium replicate.

To validate that this simulation pipeline generates realistic Visium-like data, we compared ctSVG detection results on the Xenium-derived simulated datasets with results on real Visium datasets, where the Visium and Xenium datasets were generated from serial sections of a tissue block (see Supplementary Methods S1.5 for details). The results in Fig. 3b show that when we use the overlap of the top detected ctSVGs between two datasets to measure their similarity, the similarity between our simulated datasets and real Visium datasets was close to the similarity between biological replicates. This implies that the technical variation introduced by our simulation process was within a reasonable range.

Across all five benchmarked methods, detection accuracy reduced substantially in *realistic simulations* compared to *idealized simulations*, with average EP for most methods dropping from above 80% to below 20%. As shown in Fig. 2, CELINA showed comparatively strong and consistent performance across the realistic datasets, with an average EP of 16%. Notably, despite its instability in *idealized simulations*, spVC attained the highest detection accuracy in two of the three realistic datasets, with an EP value of 41% in breast tumor and 24% in ovarian cancer. CTSV followed closely in these two tissues, with EP values of 37% and 20%, respectively. However, the performances of both spVC and CTSV worsened notably and exhibited nearly no detection accuracy on the lymph node dataset, which is the dataset with the weakest spatial pattern.

Although the statistical calibration of all methods remained within the range observed in *idealized simulations*, STANCE, C-SIDE, CTSV, and spVC all displayed severe inflation of false discoveries in *realistic simulations*. Their average FDPs reached 98%, 81%, 73%, and 72%, respectively, at a target FDR of 5%. CELINA was relatively better, but still achieved an average FDP of 41%. These results indicate that while statistical calibration was largely preserved, the ability of all methods to correctly rank cell-type-specific spatial signals deteriorated substantially under realistic tissue complexity and weaker, noisier ctSVG signals.

### 2.4. Diagnosing detection performance deterioration using *decomposed simulations*

The poor detection accuracy and false discovery control of all methods in the *realistic simulations* motivated us to investigate which components of the *realistic simulations* cause these methods to fail. To better understand the substantial gap between *idealized simulations* and *realistic simulations*, we decomposed their differences into interpretable components and quantified their impact on detection performance. We grouped these components into two major aspects of realness—tissue structure and gene expressions. The complexity of tissue structure includes three realness components: (i) the number of cell types and their proportions in tissues; (ii) cell and spot locations; (iii) cell-type composition patterns. The complexity in gene expression also includes three realness components: (iv) distribution of null genes; (v) Visium capture efficiency; (vi) intra-cell-type spatial patterns.

We conducted the *decomposed simulations* using the *idealized simulation* Scenario 1 as a starting point. We added different realness components individually or sequentially to generate simulated datasets with varying levels of realness, bridging the gap between the *idealized simulations* and the *realistic simulations*. On the one hand, we added the following components individually to the baseline dataset: (i) the number of cell types and their proportions in tissues, (ii) cell and spot locations, and (iv) distribution of null genes, to directly quantify their impacts on method performance. On the other hand, we added the following components sequentially to the baseline dataset: (iii) realistic cell-type composition patterns, (v) Visium capture efficiency, and (vi) intra-cell-type spatial patterns, as they depend on other components. For example, component (iii), realistic cell-type composition patterns, depends on both realistic (i) cell types and (ii) locations. The effects of these components were quantified using comparisons between combinations with and without the component. Method performances in all the components and combinations of components included in the *decomposed simulations* are summarized in Fig. 4a.

**Fig. 4:**
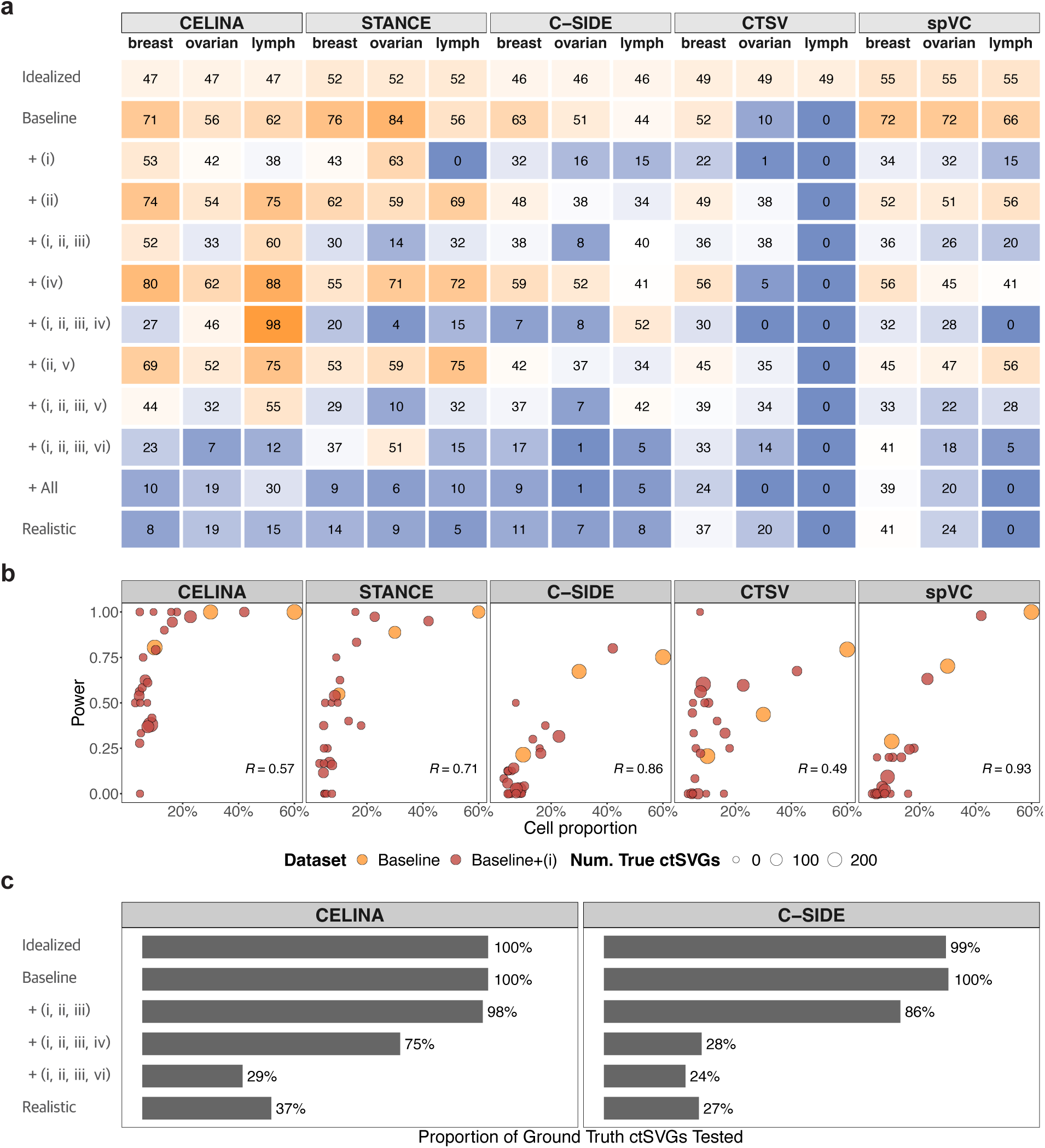
Diagnosing method performance using *decomposed simulations*. **a** Heatmap of EP (in percentage) across different simulation settings and tissue types. The simulation settings range from idealized settings (top), to decomposition settings, which introduce various tissue structure and gene expression components and their combinations, and finally to realistic settings. **b** Relationship between cell-type proportion and detection power for Scenario 1 of the *idealized simulations* and the “+ (i)” setting of the *decomposed simulations*. **c** Proportion of ground-truth ctSVGs tested by CELINA and C-SIDE in different simulation settings.

#### 2.4.1. Most methods maintained strong performance in the adapted idealized simulations, but CTSV was sensitive to signal prevalence

In the *decomposed simulations*, we first generated baseline datasets for the three tissue types—breast tumor, ovarian cancer, lymph node—based on Scenario 1 in Results 2.2. We made several minor adaptations to the simulation setting to better integrate the decomposition framework. For example, the number of genes and true ctSVGs were based on the corresponding real datasets. Full details of these adaptations are provided in Supplementary Methods S1.11. This adapted dataset served as the baseline for decomposition analysis, in which we generated nine *decomposed simulations* for each tissue type relative to the baseline.

The results in Fig. 4 show that, as the baseline retained the idealized assumptions, most methods maintained a strong detection performance. However, CTSV deteriorated sharply: its average EP for ovarian cancer and lymph node datasets dropped from 48% and 50% in the *idealized simulations* to 10% and 0% in the baseline of the *decomposed simulations*. The poor performance of CTSV in these datasets was driven by its inflated false discoveries. In the baseline, CTSV reported 5.6% and 3.9% of the genes as discoveries in the ovarian cancer and lymph node datasets, with the ground truth proportions in those datasets being only 0.6% and 0.1%, respectively. At a target FDR of 0.05, its actual FDP was 94% and 99% in those two datasets, respectively. This severe lack of FDR control is consistent with CTSV’s anti-conservative statistical calibration described in Results 2.2. These results suggest that as truth ctSVGs become increasingly rare, CTSV struggles to distinguish true signals from spurious patterns arising from non-ctSVGs.

#### 2.4.2. All methods showed substantial loss of accuracy when real tissue structures were introduced

We examined the effects of real tissue structures, including (i) realistic cell-type numbers and proportions, (ii) realistic cell and spot locations, and (iii) realistic cell-type composition patterns.

In the “+ (i)” setting of the decomposition analysis, we introduced realistic cell-type numbers and proportions in order to match the diversity of cell types in real tissues. In the “+ (ii)” setting, we added realistic cell and spot locations to the baseline data, replacing the idealized uniform cell density with the heterogeneous spatial layouts found in real datasets, including regions of high density and occasional voids. Because realistic cell-type composition patterns depend on both realistic cell-type diversity and realistic spatial location, in the “+ (i, ii, iii)” setting, we first incorporated components (i) and (ii) and then assigned each cell location to the cell type occupying that location in the real tissue (Fig. 1d). In total, we considered three additional “decomposed simulation” settings for real tissue structures: baseline “+ (i)”, “+ (ii)”, and “+ (i, ii, iii)”.

The results in Fig. 4a show that, when comparing the baseline with the “+ (i)” setting, the EP of all methods reduced substantially across tissue types, with a 9%–56% reduction in EP after adding component (i). To investigate why realistic cell-type numbers and proportions led to the reduction in detection power, we examined the detection power across cell types. The results in Fig. 4b show a strong positive correlation between detection power and cell-type proportions. Moreover, all methods achieved comparable power for cell types with similar abundance, regardless of whether the dataset came from the baseline or the “+ (i)” setting. These results indicate that the detection power is negatively affected by low cell type abundance, and the low power in rare cell types leads to the reduced overall detection power. Notably, random-effect models, CELINA and STANCE, were more robust to low cell-type proportions, maintaining consistently higher detection power than fixed-effect models (C-SIDE, CTSV, spVC) among minority cell types comprising less than 10% of total cells.

Comparing the “+ (i)” and “+ (i, ii, iii)” settings, the EP for most methods has not changed significantly, indicating that components (ii) and (iii) had a relatively low impact on detection accuracy. However, STANCE is an exception as its EP declined drastically in breast and ovarian cancer datasets. Its FDP increased from 54% in “+ (i)” to 84% in “+ (i, ii, iii)” (Supplementary Fig. S2). As discussed in Results 2.2, STANCE has inflated Type I errors for cell-type marker genes as it mis-attributes spatial variation driven by non-uniform cell-type composition to ctSVG signals. In “+ (i, ii, iii)” which incorporates realistic locations and composition patterns, the type I error of STANCE for marker genes further increased to 47% on the heterogeneous ovarian cancer tissue (Fig. 3)

#### 2.4.3. Some methods showed method-specific vulnerability towards realistic gene expression patterns

Finally, we examined the effects of the realistic gene expression patterns, including (iv) distribution of null genes, (v) Visium capture efficiency, and (vi) intra-cell-type spatial patterns. These components modify the gene expression levels of the simulated datasets to mimic biological reality. To add component (iv) to the *decomposed simulation* settings, we introduced realistic null gene expressions. Different from *idealized simulations* where all non-marker and non-ctSVG genes share identical mean expression across cell types and spatial locations, we used scDesign3 to fit the expression distributions of null genes within each cell type and generated their counts from the fitted models. To add component (v) to the *decomposed simulation* settings, we incorporated the transcript loss observed in real Visium datasets by randomly removing about 50% of the transcripts. The removal proportion was calibrated using the proportion of transcripts that fall within the Visium capture discs. Finally, to add component (vi) to the *decomposed simulation* settings, we introduced realistic intra-cell-type spatial patterns. In *idealized simulations*, ctSVGs follow a simple predefined spatial pattern (e.g., patterns illustrated in Fig. 1b). Here, we used scDesign3 to fit the spatial expression patterns for the selected ground truth ctSVGs and generated simulated gene expressions from the fitted model. Due to the same reason as illustrated in the previous section that some components are dependent on others, we considered six additional “decomposed simulation” settings: baseline “+ (iv)”, “+ (i, ii, iii, iv)”, “+ (ii, v)”, “+ (i, ii, iii, v)”, “+ (i, ii, iii, vi)”, and “+ All”.

The results in Fig. 4 show that components (iv) and (vi) affect detection accuracy for some methods, whereas component (v) had minimal effect on all methods. Specifically, compared to “+ (i, ii, iii)”, STANCE’s FDP inflated from an average of 84% to 98% under “+ (i, ii, iii, iv)” (Supplementary Fig. S2). The effect of component (iv) on STANCE is similar to that of component (iii), which likely arises from its limited ability to separate spatial variation driven by non-uniform cell-type composition and intra-cell-type spatial pattern (see Results 2.2). Component (iv) assigns distinct mean expression levels for non-ctSVGs across cell types, introducing some cell-type marker genes. As the cell-type composition is spatially non-uniform, these marker genes exhibit spatial variation that STANCE often mis-identifies as ctSVGs.

Compared with “+ (i, ii, iii)”, the detection accuracy of C-SIDE and CELINA dropped substantially under “+ (i, ii, iii, iv)”, indicating a substantial effect of component (vi). This effect likely arises from the quality-control filters implemented by these two methods, which excluded the majority of ground truth ctSVGs in the settings with realistic spatial patterns. The filtering steps were only briefly noted in their papers and were designed to remove genes with low expressions. As shown in Fig. 4c, in the *idealized simulation* Scenario 1, true ctSVGs have relatively high expressions by design, and both CELINA and C-SIDE tested nearly all ctSVGs. In “+ (i, ii, iii)”, CELINA and C-SIDE only filtered out 2% and 14% of true ctSVGs, respectively. However, in “+ (i, ii, iii, vi)”, where realistic spatial patterns were added, the proportion of filtered out true ctSVGs increased sharply to 71% for CELINA and 76% for C-SIDE. This substantial loss of true ctSVGs is likely due to weaker spatial signals under realistic patterns. As a result, the attainable detection accuracy of CELINA and C-SIDE became fundamentally limited by these quality-control filters.

### 2.5. Computational scalability in time and memory

We next evaluated each method’s time and space scalability under varying numbers of spots and genes, using simulated datasets generated from the *idealized simulation* setting: Scenario 1 and Hotspot 1 (see Fig. 1b). To assess how scalability is influenced by the number of spots, we fixed the number of genes at 100 and varied the number of spots from 100 to 40,000. To assess how scalability is influenced by the number of genes, we fixed the number of spots at 900 and varied the number of genes from 100 to 20,000. All these simulations were run on standard virtual machines equipped with 16 CPU cores and 245 GB memory (see Supplementary Methods S1.8 for details).

Because several methods showed convergence issues in the *realistic simulations*, we also evaluated the scalability using *decomposed simulations*, where the numbers of spots and genes were fixed but the tissue structure and gene expression complexity were increased. *Decomposed simulation* results were normalized for comparability using CPU time, defined as the product of the number of CPU cores and elapsed time.

spVC and CTSV both demonstrated efficient memory usage and successfully completed all scalability experiments (Fig. 5a). spVC exhibited the best overall efficiency in both runtime and memory. However, its runtime fluctuated substantially across decomposition experiments, increasing by up to two orders of magnitude relative to the *idealized simulation* once all realness components were added (Fig. 5b). CTSV achieved similar memory efficiency as spVC, yet its runtime exceeded two hours for datasets with over 10,000 genes, consistent with prior reports that CTSV is computationally prohibitive for large-scale analyses. C-SIDE maintained runtime below five minutes across all scalability experiments and was stable in the *decomposed simulations*, but it failed on datasets with over 10,000 genes because its quality-control filters excluded all genes from testing.

**Fig. 5:**
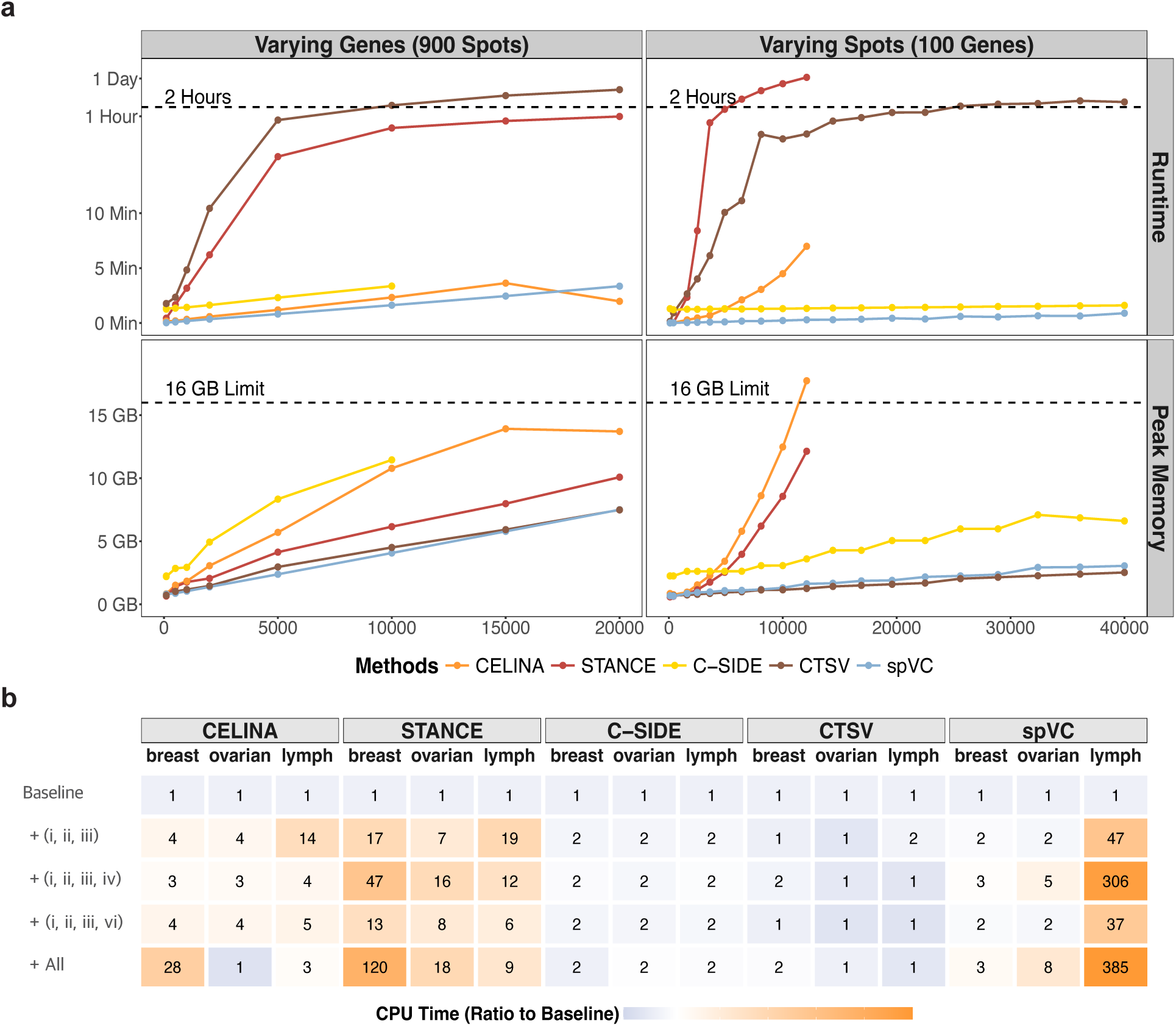
Benchmarking runtime and memory scalability across varying dataset sizes. **a** Runtime and peak memory usage for each method as the number of genes (900-spot setting) or spots (100-gene setting) increases, evaluated on a 16-core, 245-GB virtual machine. **b** Heatmap showing relative CPU time in *decomposed simulations*, expressed as the ratio to each method’s CPU time in the baseline setting.

Random-effect methods, STANCE and CELINA, encountered memory constraints as the number of spots increased, with both exceeding 245 GB limit once datasets reached 12,100 spots. STANCE’s runtime grew rapidly due to the slow EM algorithm convergence, producing frequent non-convergence warnings beyond 400 spots and exceeding two hours for datasets with more than 4,900 spots. CELINA maintained runtimes below 10 minutes for datasets that did not reach the memory limit, though runtime also rose sharply with the number of spots. Overall, both runtime and memory requirements scaled unfavorably for STANCE, while memory exhaustion remained the primary bottleneck for CELINA. Both methods also showed unstable runtime in the *decomposed simulations*: despite the fixed numbers of spots and genes, STANCE’s CPU time increased by up to 120-fold and CELINA by up to 28-fold once all realness components were added (Fig. 5b).

## 3. Discussion

In this work, we conducted the first-ever comprehensive benchmark study of all existing ctSVG detection methods. To ensure that our evaluation reflects real-world performance of the methods, we simulated biologically realistic datasets using scDesign3. Our results show that although current ctSVG detection methods perform reasonably well under idealized settings, they tend to struggle under realistic settings. Across tissue types and disease states, they identified fewer than 20% of ground-truth ctSVGs among their top-ranked discoveries. Moreover, none of the methods controlled false discoveries reliably. The FDP was already inflated in *idealized simulations*, and this inflation substantially worsened in *realistic simulations*, indicating the limited ability of existing methods to provide reliable discoveries. To understand why these failures occur in *realistic simulations*, we proposed a new decomposition framework, which decomposes the realness of the *realistic simulations* into interpretable components. This framework enables us to quantify how specific aspects of tissue structure and gene expression complexity affect detection accuracy and error control. For example, we identified that the quality-control filters in CELINA and C-SIDE restricted their detection power when true signals are comparatively weak, while STANCE yielded inflated false discoveries because it mis-identifies cell-type marker genes as ctSVGs.

In addition to detection accuracy on simulated datasets, we also compared these methods in terms of their computational scalability and overall usability. We then synthesized all benchmarking results into a practical user guide that highlights practical considerations for applying existing ctSVG methods in real data analysis. A detailed comparison of input requirements and evaluation metrics is shown in Fig. 6. No single method dominates the others across all evaluation criteria. For example, spVC achieved the best detection performance, but its detection accuracy was unstable and dropped to near-zero accuracy on the realistic dataset with the weakest signal; CELINA achieved relatively stable detection performance but can become memory-prohibitive on datasets consisting of more than 10,000 spots. Given these trade-offs, we recommend that users select methods by aligning their dataset characteristics and analysis priorities with the strengths and limitations of the methods summarized in the user guide.

**Fig. 6:**
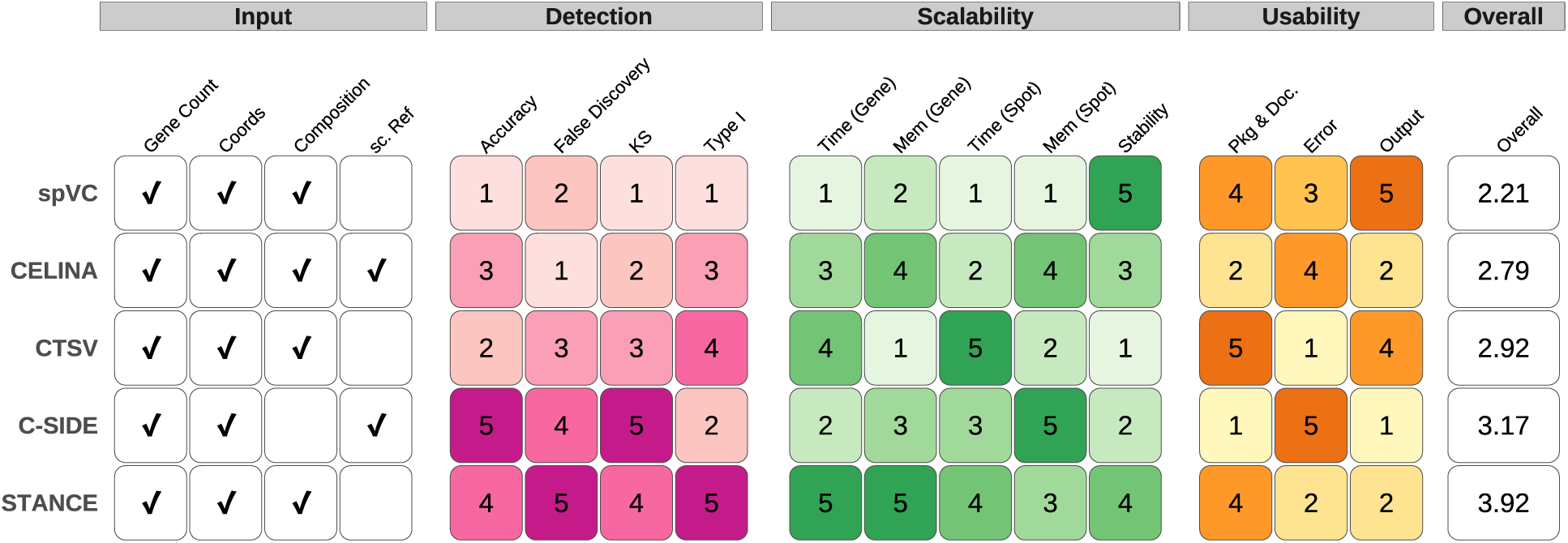
Summary of the benchmarked methods, including the required inputs for each method (the leftmost column), and their performance in terms of detection accuracy, scalability and usability (the middle three columns). In the performance heatmaps showing the performance of the methods, the number indicates each method’s rank (smaller is better) among all five methods for each metric. The rightmost column shows the overall average ranking of the five methods across all metrics (smaller is better).

Beyond guiding method selection, our *decomposed simulations* highlight three key challenges that future ctSVG methods should address. First, we observed that detection power is negatively correlated with cell-type abundance for all methods. Cell types with proportions below 10% are very common in real tissues, and ctSVGs from these cell types are as biologically important as those from more abundant cell types. However, for most existing methods, the detection power for ctSVGs in rare cell types is very low, and for some methods, the power is nearly zero. Improving the detection power for minority cell types is thus a central challenge for future development of ctSVG detection methods. Second, gene-level quality-control filtering is widely used to exclude noisy, lowly expressed genes and accelerate computation. However, the filters implemented by CELINA and C-SIDE often remove most true ctSVGs and thus place a hard limit on the attainable accuracy in realistic tissues, where signals are weaker and less consistent. Improving these filters is therefore another key step for enhancing the detection power. Third, scalability also represents a critical bottleneck. Although CELINA and CTSV show strong detection performance, both face substantial memory or runtime demands as the number of spots or genes increases, limiting their applicability to finer resolution datasets containing tens to hundreds of thousands of pixels. Continued development of computationally efficient, statistically calibrated, and biologically informed ctSVG methods will be necessary to unlock the full potential of spatial transcriptomics with increasing resolution and scale.

The decomposable diagnostic framework developed in this work also extends naturally to other omics data analysis, helping facilitate benchmarking and enabling method developers to more accurately identify weaknesses in their methods. For example, although computational tools for detecting spatially variable chromatin accessibility or protein abundance remain limited, spatial ATAC-seq and spatial proteomics face many of the same modeling challenges observed in spatial transcriptomics, including large numbers of cell types, substantial technical and biological noise, and weaker or less consistent spatial structure in real tissues. A decomposition-based strategy can help identify which aspects of biological and technical complexity most undermine performance. More broadly, the same principles apply to single-cell trajectory inference, where many methods are evaluated using simulated datasets built on over-simplified assumptions about lineage structure and expression noise. As reference-based simulation approaches become increasingly common for generating realistic single-cell and spatial data, a systematic decomposition of realness complexity can provide a principled way to assess robustness, diagnose failure modes, and guide the development of more resilient computational methods across modalities.

Several limitations of this study are worth noting. First, our *realistic simulations* were designed to mimic the widely used Visium platform, and we did not evaluate how it generalizes to other spot-resolution technologies such as ST [24], Slide-seq [25], or Stereo-seq [26]. Extending the benchmark to these platforms will be important given their distinct spatial resolutions and capture characteristics. Second, although our real datasets span multiple tissue types and disease states, they may not capture the full biological diversity encountered in practice. Additional tissues may reveal further method-specific behaviors. Finally, both our *idealized* and *realistic simulations* assume that spot-level cell-type compositions are known, mirroring the assumptions made by all existing ctSVG methods. In practice, these compositions must be estimated using deconvolution methods [27, 28]. How uncertainty in these estimates influences ctSVG detection has not been examined. Assessing the robustness of existing methods to imperfect or biased cell-type decomposition remains an important direction for our future work.

