## Supplementary File for "Benchmarking Cell-Type-Specific Spatially Variable Gene Detection Methods Using a Realistic and Decomposable Simulation Framework"

### S1. Supplementary Methods

#### S1.1. Computational methods for ctSVGs detection

In this subsection, we provide a brief introduction to the five benchmarked method. All of them model gene expression as a function of cell-type constant effects, intra-cell-type spatial variation, and a library size component. Based on how they model the spatial variations of the ctSVGs, we group these methods into two categories: fixed-effect models and random-effect models, which differ in three major aspects. First, fixed-effect methods model raw gene counts, whereas random-effect methods operate on normalized expression values, reflecting the distributional assumption required by their Gaussian process formulation. Second, as illustrated in Fig. S1, fixed-effect models include all cell-type-specific effects and incorporate library size explicitly as an exposure, while random-effect models omit some cell-type-specific effects and account for library size implicitly through normalization. Third, although all methods ultimately report p-values for each gene-cell-type pair, they differ in how those p-values were generated: fixed-effect methods fit a single model per gene and test multiple gene-cell-type effects within that model, whereas random-effect methods fit an independent model for every gene-cell-type pair.

Beyond modeling of the spatial variation, the methods also differ in their quality-control filters, specification of spatial effects, and approaches for computing and adjusting p-values. Although those modeling choices are often understated, they have substantial implications for statistical calibration and detection accuracy as examined in Results section. Below, we provide additional detail on these aspects.

**CELINA**. Prior to testing, CELINA implements similar gene-level quality control filters as C-SIDE. Specifically, it only conducts test for genes with normalized expression above a user-specified threshold and have mean expression in target cell type over half of the its highest expression among other cell types. For all remaining gene-cell-type pairs, CELINA fits 11 random-effect models, each modeling the target cell's intra-cell-type spatial variation with covariance matrix constructed based on different kernel including 5 Gaussian, 5 Matérn and 1 spline families. As it conducts one Score test for each kernel, it combines those results through Cauchy combination test, yielding a single p-value per gene-cell-type pair. While CELINA doesn't perform multiple testing adjustments, we acquire global B-H adjustment for all gene-cell-type pairs in each dataset.

**STANCE**. STANCE uses a two-stage strategy to identify ctSVGs. In the first stage, STANCE tests all genes to identify utSVGs—genes that exhibit any spatial expression pattern. In the second stage, only the identified utSVGs proceed to a follow-up test to identify ctSVGs, which have spatial patterns specific to certain cell types. Before stage one, STANCE performs a quality-control step which filters out genes with normalized expression below the threshold. To model intra-cell-type spatial effect, STANCE constructs covariance matrix with only one Gaussian kernel and thus produces a p-value for each gene-cell-type pair directly through Score test. Similar as CELINA, STANCE also doesn't offer multiple testing adjustment, and we handle it as above described in CELINA.

**CTSV**. CTSV does not have a filtering step and tests all possible gene-cell-type pairs. It models the intra-cell-type spatial effect of each cell type present in the tissue with two sets of five kernel functions built on x and y coordinates respectively, including 2 Gaussian kernel, 2 periodic kernels and 1 linear kernel. To produce a single p-value for each gene-cell-type pair, it first combines each set of kernel results through Cauchy Combination test, and CTSV claims intra-cell-type spatial effect as long as one of the two null hypothesis is rejected and thus implicitly taking the minimum of the two p-values. Instead of directly apply B-H on the final p-values, CTSV applies B-H across p-values acquired for x and y coordinates for all gene-cell-type pairs and then take the minimum of the x and y coordinate adjusted p-value as the final adjusted p-value.

**C-SIDE**. C-SIDE have similar gene level quality control mechanism as CELINA. C-SIDE models the intra-cell-type spatial effect for each cell-type with a set of thin plate spline basis functions, the number of which can be specified by users. C-SIDE conduct a Wald test for each of the basis functions. In order to produce a single p-value for each gene-cell-type pair, it takes the minimum of Bonferroni corrected p-values of those basis functions. Subsequently, C-SIDE conducts B-H multiple testing adjustment within each cell type.

**spVC**. spVC filters out any genes has lower than 100 count across all spots. spVC models the intra-cell-type spatial effect for each cell-type with basis functions from bi-variate penalized smoothing spline over triangulation (BPST). Although it has multiple basis functions representing one gene-cell-type spatial effect, it conducts a Wald-like test to obtain one calibrated p-value for each smoothing component and thus avoids combining multiple p-values. Similarly as STANCE and CELINA, spVC doesn't have built in multiple testing adjustment, we conduct B-H adjustment across gene-cell-type pairs in each dataset.

### S1.2. Idealized simulation framework

The idealized simulations were reproduced following the designs of STANCE and CELINA. Each simulation first generated cell-level $(x, y)$ coordinates, cell-type labels, and gene expression counts.

Cells were then aggregated into spots arranged on equal-sized square grids. Each square corresponded to a spot, and spot-level measurements were obtained by aggregating the cells falling within that square: gene counts were summed across cells, spot coordinates were computed as the mean of cell coordinates, and cell-type compositions were defined as the proportion of cell types among those cells. Since spot-level data were direct aggregation of cell-level data, we describe only the cell-level data generation below.

For all idealized simulations, cell locations were assumed to follow Poisson random point process on the unit square. Locations were generated using the `rpoispp()` function in the *spatstat* package with specified expected cell counts and a unit-square window. After locations were generated, cell types were assigned according to the cell-type composition assumptions for each scenario.

- For scenario 1, cell-type composition was uniform across the tissue, with proportions of 10%, 30% and 60% for the three cell types. Cell types were therefore sampled independently for each cell according to those proportions.

- For scenario 2, composition varied across spatial domains. Two circular regions were defined by randomly selecting two generated cell locations as centers an drawing radii from a unifrom distribution between $0.1$ and $0.5$. Together, the two circles and the area outside them formed three spatial domains. Each cell type was present in only two of the three domains, with 50% abundance within each domain it occupied. Cell types were then sampled within each domain based on these domain-specific proportions.

- For scenarios 3-6, cell-type composition varied across four rectangular spatial domains created by evenly partitioning the tissue along the x-axis, while each cell type contributed equallyto the total number of cells. Scenario 3 had minimal overlap: each of the four cell types was restricted to its own domain. In scenarios 4-6, each cell type appeared in three domains with decreasing dominance in their home domain. Specifically, each cell type occupied its home domain with proportions of $90\%$, $50\%$, and $33\%$ (scenarios 4-6), and the remaining proportions was distributed evenly across the non-dominant domains ($5\%$, $25\%$ and $33\%$ per domain, respectively).

For all these scenarios, the spatial expression patterns of the ground-truth ctSVGs come from one of the following:

- **Hotspot 1**. A circular domain was centered at a randomly selected cell location, with its radius drawn from a uniform distribution between $0.2$ and $0.4$. Each ground-truth ctSVGs exhibited a specific fold change between the gene expressions of the cells inside the domain and those outside it. The fold change for each ctSVG was randomly chosen from $\{4, 2, 0.5, 0.25\}$ with equal probability.

- **Hotspot 2**. For each ground truth ctSVG, we define a circular domain whose center was the median coordinate of all cells belonging to that ctSVG's cell type. The radius was then chosen so that a preset proportion of those cells fell inside the domain. Half of the ground-truth ctSVGs were up-regulated within the domain: the expressions of the cells in the right half of the domain had a fold change of 2 relative to cells outside the domain, and the cells within the left half of the domain had a fold change of 2 relative to those in the right half. For the other half of down-regulated ctSVGs, the corresponding fold changes were 0.4 (right vs. outside) and 0.5 (left vs. right).

- **Streak**. For each ground-truth ctSVG, a rectangular domain was defined by slicing from the tissue along the y-axis. The height of this domain was chosen so that a preset proportion of cells belonging to that ctSVG's cell type fell inside the domain. Half of the ground-truth ctSVGs were up-regulated within the domain with a fold change of 1.5 between cells inside and outside the domain; the other half were down-regulated with a fold change of

- **Gradient**. For each ground-truth ctSVG, we induced a gradient pattern by reordering its expression values according to the cells' y-coordinates. Specifically, expressions values of the ctSVG were first generated for all cells from a homogeneous distribution without spatial pattern. Then, a preset proportion of the cells was randomly sampled, and their expression values were re-ordered so that the ordering of their expression matched the ordering of their y-coordinates, either in ascending or descending direction. Half of the ground-truth ctSVGs were assigned an increasing gradient of expression from the top to the bottom of the y-axis, while the other half were assigned a decreasing gradient.

We then selected a predefined number of marker genes and ground-truth ctSVGs from a total of 1,000 genes. Each cell type was assigned a fixed number of marker genes, which were up- or down-regulated only within the designated cell type. Scenario 1 assumed that each cell type had its own set of ctSVGs that did not overlap with marker genes or with each other, and all ctSVGs followed the Hotspot 1 pattern. Scenario 2 was a null setting with marker genes only and no ctSVGs. For scenarios 3–6, we generated four datasets per scenario: one null dataset with marker genes only, and three non-null datasets corresponding to the three ctSVG spatial patterns. In non-null datasets, only one randomly selected cell type contained ctSVGs. All remaining genes were null genes with constant mean expression across both cell types and spatial locations.

For intra-cell-type patterns other than Gradient, we simulated the cell level gene expression counts based on the above defined mean expression for each gene through negative binomial distribution `nbnomial()`. The size parameter for negative binomial distribution is a preset constant for all genes, 0.7 and 1.5 were used to generate separate sets of scenario 1 and scenario 2 simulations, and only 0.7 is used for all scenario 3-6 simulations.

### S1.3. Real datasets used in this study

We analyzed three publicly available spatial transcriptomics datasets from 10x Genomics.

1. Breast tumor (Xenium + Visium).Two Xenium *in situ* datasets and one Visium dataset were generated from serial sections of the same human breast tumor [1]. These data are available

at https://www.10xgenomics.com/products/xenium-in-situ/preview-dataset-human-breast. Each Xenium replicate contains transcript-level counts and transcript coordinates. The Visium dataset contains spot-level gene counts. We used the associated H&E images to align the three serial sections via affine transformations exported from Xenium Explorer. For benchmarking, we used spot-level Visium data for concordance analyses and Xenium replicate 1 to generate the *realistic simulations* described in Section S1.4.

2. Ovarian cancer (Xenium). The FFPE human ovarian cancer Xenium dataset (available at https://www.10xgenomics.com/datasets/xenium-prime-ffpe-human-ovarian-cancer) contains transcript-level counts, transcript coordinates, cell–transcript assignments, and an accompanying H&E image. We used this dataset to construct an additional realistic simulation setting.

3. Lymph node (Xenium).The human lymph node Xenium dataset (available at https://www.10xgenomics.com/datasets/preview-data-xenium-prime-gene-expression) provides transcript-level expression, transcript coordinates, cell–transcript mappings, cell type annotations, and an H&E image.

For all Xenium datasets, we used the cell–transcript assignments and the cell type annotations provided by 10x Genomics. To our knowledge, no peer-revised publication is currently associated with the ovarian cancer and lymph node dataset. We therefore cite them directly from 10x Genomics.

#### S1.4. Realistic simulation framework

The simulation of realistic datasets involved four main steps: (1) fitting cell-level counts with scDesign3, (2) selecting ground-truth ctSVGs, (3) generating new cell-level data, and (4) aggregating simulated cells into Visium-like pseudo-spots. We based all simulations on 10x Genomics Xenium subcellular-resolution datasets to ensure realistic cell-type-specific spatial pattern and accurate cell-type annotations. Simulating counts at the cell level avoids reliance on (i) cell type deconvolution models that infer spot-level cell-type composition and (ii) complex spot-level models that jointly estimate cell-type constant effects and spatial effects.

Let $C_{ig}$ denote the observed count of gene $g$ in cell $i$, where $i \in \{1, \dots, N\}$ indexes cells and $g \in \{1, \dots, G\}$ indexes genes. Each cell $i$ belongs to cell type $k(i)$ with spatial coordinates $(x_i, y_i)$. For each cell type k with at least $n_k \geq 100$ cells, scDesign3 models expression as

$$C_{ig} \sim \mathrm{NB}(\mu_{ig}, \theta_{kg}), \quad \log \mu_{ig} = f_{kg}(x_i, y_i), \tag{S1}$$

where both the mean $\mu_{ig}$ and dispersion $\theta_{kg}$ are estimated. The smooth spatial function $f_{kg}(\cdot)$ is modeled by a Gaussian process over cell coordinates $(x_i, y_i)$ with kernel bandwidth 50. For each gene $g$ within cell type $k$, the fraction of deviation explained by the marginal model,

$$R^2_{kg} = 1 - \frac{\mathrm{Deviation}_{kg}}{\mathrm{Null\ Deviation}_{kg}}, \tag{S2}$$

were recorded to guide ctSVG selection. Gene-gene dependence within each cell type was modeled using a Gaussian copula over the fitted marginals.

To identify candidate ctSVGs, we computed Moran's Index using function `FindSpatiallyVariableFeatures()` from package *Seurat* and retained the top 200 genes per cell type. We excluded genes expressed in $< 1\%$ or $< 100$ cells of corresponding cell type so that the spatial pattern is genuine instead of artificially high Moran's Index and deviation explained due to few non-zero expressions. We also restricted selection to cell types collectively representing $\geq 90\%$ of all cells to ensure stability in downstream analysis. The remaining gene were ranked by $R^2_{kg}$, and the top $N_{\text{ctSVG}}$ were visually confirmed to display interpretable spatial structure (e.g., $N_{svg} = 20$ in Xenium lymph-node dataset).

The estimated mean was modified to define null versus ctSVGs. Specifically, we preserved fitted parameters for genes selected as ground truth ctSVGs while replacing fitted means of null genes to the average fitted mean for that cell type to eliminate any intra-cell-type spatial pattern.

$$\tilde{\mu}_{ig} = \begin{cases} \frac{1}{n_k} \sum_{i' \in \mathcal{I}_k} \mu_{i'g}, & \text{if } g \in \text{null genes}, \\ \mu_{ig}, & \text{if } g \in \text{ctSVGs}. \end{cases} \tag{S3}$$

New cell-level counts $\tilde{C}_{ig}$ were simulated using `simu()` in `scDesign3` with the above adjusted mean parameters, scDesign3 estimated dispersion $\theta_{kg}$ and gene-gene correlation structure.

To simulate Visium-like spot-level data, we generated a grid of 55 $\mu m$-diameter capture discs with 100 $\mu m$ spacing following the 10x Visium capture specification [2]. Each transcript was assigned to spot if its coordinates fell within the capture disc. We defined a cell-to-spot matrix $A \in \mathbb{R}^{N \times M}$, where $A_{im}$ is the proportion of assigned transcripts from cell $i$ assigned to spot $m$. We also defined a drop matrix $D \in \mathbb{R}^{N \times G}$, where $D_{ig}$ represents the proportion of transcripts of gene $g$ from cell $i$ that fall outside all capture regions. The pseudo-spot count matrix $S \in \mathbb{R}^{M \times G}$ was then computed as

$$S_{mg} = \sum_{i=1}^{N} A_{im}(1 - D_{ig})\tilde{C}_{ig}. \tag{S4}$$

Finally, the cell-type decomposition matrix $P \in \mathbb{R}^{M \times K}$ was obtained by mapping each cell in A to its annotated type $k(i)$ and normalizing the result by each spot so that the column add up to 1.

### S1.5. Validation of realistic simulations with Visium data

The above simulation pipeline extends scDesign3 by generating pseudo-spot-level data from subcellular inputs, rather than simulating data which has the same resolution as the input. Therefore, it is necessary to verify that the simulated data from scDesign3 resembles real Visium data, and that the pseudo-spot aggregation strategy does not alter the behavior of ctSVG detection methods beyond a reasonable range. To this end, we used a 10x Genomics breast cancer dataset in which multiple serial sections of the same tissue sample were sequenced with both the Visium and Xenium platforms [1].

We used two Xenium replicates and one Visium replicate. The H&E images of all three serial sections were co-registered, and we selected the overlapping region shared across all replicates. We then generated ten synthetic pseudo-spot datasets from Xenium replicate 1 following steps 1, 3 and 4 of our simulation pipeline — fitting each cell type with scDesign3 and aggregating simulated cells into pseudo-spots. Because real Visium spot coordinates were available on this dataset, we directly adopted those coordinates to define capture spots of 55 $\mu m$-diameter [2]. With the same defined capture spots, we turned Xenium replicate 1 and 2 into pseudo-spot data by aggregating the sequenced transcripts within the bound of each capture spot.

We quantified similarity between two datasets by measuring the proportion of overlapping ctSVG among their top $N$ discoveries for each method, where $N$ ranged from $1$ to $200$. We focus on two similarities in our comparisons. The first similarity that we assessed was the similarity between the simulated dataset and the real Visium dataset. We then compared this similarity with two baseline similarities: (1) the similarity between Xenium replicates 1 and 2, which reflects the agreement between biological replication, and (2)the similarity between the real Xenium and Visium datasets, which captures the combined effects of biological replication and platform difference. Our results show that the similarity between the synthetic dataset and the real Visium dataset was nearly identical to baseline similarity (1), indicating that the discrepancy introduced by our pseudo-spot simulation process fall well within the natural combined effects of biological replicates and platform difference. When compared against the baseline similarity (1), the similarity between the synthetic dataset and the real Visium dataset remained on the same scale.

The second similarity that we assessed was the similarity between two synthetic datasets generated by our pseudo-spot simulation based on scDesign3, which reflects the reproducibility of the simulations process. Specifically, we compared the similarity between two synthetic datasets generated from Xenium replicate 1 with the similarity between one synthetic dataset and the real Xenium replicate 1. The results show that these two similarities were highly comparable. This further confirms that the our simulation framework generates reliable and reproducible pseudo-spot datasets.

### S1.6. Decomposed simulation framework

Detection performance deteriorated substantially on realistic datasets compared to idealized simulations. To investigate this discrepancy, we developed a diagnostic framework that decomposes the realness of realistic datasets into interpretable components. Starting from idealized simulation Scenario 1 with the Hotspot 1 pattern, we added individual realness components or representative combinations of them one at a time.

Before introducing any realness components, we made four adaptations to ensure compatibility with realistic settings and to avoid any potential confounding. First, instead of simulating 1,000 genes in idealized simulation, here we matched the number of expressed genes in each Xenium dataset and randomly selected 50 marker genes per cell type from this gene set. To stabilize the ground-truth signal across decomposition experiments, we used the selected ground truth ctSVGs in each realistic dataset and sampled an equal number for each cell type, removing duplicates when necessary. Second, all ctSVGs were assigned a uniform fold change of 4 within the spatially varying domain, replacing the mixture of fold changes used in idealized simulations. Third, we adjusted the number of simulated cells and spots to match the observed cell-to-spot ratio in the real tissue. Finally, cell coordinates were rescaled from the unit square to real $(x, y)$ coordinate range.

Realistic datasets diverge from idealized simulations along two major aspects. The first aspect concerns tissue-structure complexity, consisting of (i) the real number of cell types and their proportions, (ii) the locations of cells and spots, and (iii) the resulting cell-type composition patterns. These elements are determined directly by the real tissue rather than by scDesign3's fitted parameters or Visium-like capture assumptions. Components (i) and (ii) can be introduced individually, whereas component (iii) inherently requires both, since realistic cell-type composition patterns depend jointly on the diversity of cell types present and their spatial layout.

To incorporate (i), we replaced the three synthetic cell types with the full set of cell types observed in the Xenium dataset and randomly assigned the simulated cells to these cell types according to their empirical proportions. In addition, once the realistic cell-type numbers were incorporated, we could directly use the ground-truth ctSVGs defined in the realistic simulations, as the ground truth for each corresponding cell type. To incorporate (ii), we substituted the Poisson-generated coordinates with the actual Xenium cell locations and used the real cell-to-spot assignment matrix $A$ when aggregating cells to spots. Once both components (i) and (ii) were included, we further incorporated (iii) by directly assigning each real cell location to the cell type observed at that location in the Xenium data, thereby replacing the uniform composition patterns of the original idealized simulation with realistic cell-type composition patterns.

The second major aspect concerns the strength of expression signal and noise, which includes (iv) non-uniform mean expression for null genes, (v) Visium-like capture efficiency, and (vi) realistic intra-cell-type expression patterns. These components rely on scDesign3's fitted parameters or on our adjustment for capture efficiency. Component (iv) can be added independently, whereas component (v) requires real spatial coordinates (ii), and component (vi) requires the full set of tissue-structure components (i–iii).

To introduce (iv), we replaced the constant null-gene means and dispersion in idealized simulations with the scDesign3-fitted mean expression $\tilde{\mu}_{ig}$ for non-ctSVGs. When (i) was not included, the three simulated cell types were matched to the three most abundant real cell types, and their fitted means and dispersions were substituted accordingly; when (i) was included, we directly replaced the mean and dispersion of each null gene with its corresponding estimates from scDesign3. Component (v) was added on top of (ii) by first generating the idealized cell-level count matrix and then applying element-wise multiplication with $(1 - D)$, where $D$ represents the proportion of gene transcripts lying outside the Visium capture disc for each cell, thereby simulating transcript loss. Finally, to incorporate (vi), while requires components (i–iii) included, we replaced the ctSVG mean and dispersion parameters with the scDesign3-estimated parameters, introducing realistic intra-cell-type spatial variation in signal strength.

### S1.7. Detection performance evaluation metrics

We evaluated detection performance on three aspects: overall accuracy, false discoveries and detection power. Because STANCE, CELINA, C-SIDE and spVC all filter out a non-negligible proportion of gene-cell-type pairs before testing, they do not output p-values for all possible pairs. To make methods comparable, filtered-out pairs were treated as having an adjusted p-value of 1 when computing detection performance metrics. Detection accuracy was quantified using EP and AUPRC. EP was defined as the proportion of ground truth ctSVGs in the top $N_{\text{ctSVG}}$ pairs ranked by ascending adjusted p-value, where $N_{\text{ctSVG}}$ is the total number of ground truth ctSVGs in the dataset. To compute AUPRC, adjusted p-values were converted into scores using the negative natural logarithm and paired with binary labels indicating ctSVG status. The area under the precision-recall curve was obtained using function `pr.curve` from package *PRROC*. FDP and detection power were calculated at target FDR of 0.05.

$$\text{FDP} = \frac{\text{FP}}{\text{FP+TP}}, \quad \text{Power} = \frac{\text{TP}}{\text{FN+TP}}, \tag{S5}$$

where FP denotes the number of false positives, TP the number of true positives and FN false negatives.

To assess statistical calibration, we examined the K-S distance between the empirical p-value distribution of non-ctSVGs and the expected Uniform$(0, 1)$ distribution, as well as the realized type I error at nominal 0.05 level. Only gene-cell-type pairs for which a method produced p-values were included in the calculation of K–S distance. The K–S distance was computed using all tested non-ctSVG via `ks.test()`. The realized type I error was defined as the proportion of p-values below 0.05 among all non-ctSVGs regardless of whether correctly excluded before conducting the hypothesis test.

### S1.8. Scalability assessment in time and memory

To assess scalability with respect to spot number and gene number, we used idealized simulation Scenario 1 with the Hotspot 1 pattern as a base setting and varied the total number of genes and spots. We tested number of genes from 100 to 20,000 and number of spots from 100 to 40,000. To preserve the relative proportion of signal as dataset size increased, we set the number of marker genes and ctSVGs per cell type to fixed proportions of the total gene set, 10% and 5% respectively.

Computational resource usage was evaluated on standard virtual machines equipped with 16 AMD EPYC 24-core CPU processors and 245 GB of memory. All benchmarked methods provide built-in parallelization and were run using all 16 available cores. For each method, elapsed wall-clock time was measured using `system.time()`, and peak memory usage was monitored using `gc()`.

The decomposition experiments were not run on this standardized virtual-machine setup. Instead, they were executed on a mixture of CPU models, core counts, and memory configurations.

To make results from these heterogenous environments roughly comparable, we report CPU time, defined as the product of CPU cores used and the elapsed runtime.

### S1.9. Criteria for usability

We evaluated usability along three dimensions.

1. Accessibility. All five methods provide R packages, but documentation quality varies widely. C-SIDE offers the most complete support, with detailed function descriptions and tutorials covering multiple method variants. Some of the remaining tools such as CTSV provide limited examples and sparse guidance.

2. Stability and ease of debugging. CELINA applies stringent gene-level filters and requires a single-cell reference even when spot-level compositions are provided. Division-by-zero errors were common when CELINA's filtering removed genes, leaving some reference cells with zero total expression and causing failures during normalization. Because these filters are complex and largely invisible to users, debugging is difficult. C-SIDE applies both gene- and cell-level filtering and can return no results under certain settings, consistent with prior reports [3]. spVC requires manual selection of tissue boundaries, and poor boundary choices can lead to degenerate triangulations and failure.

3. Output clarity. C-SIDE, STANCE and CELINA produce cell-type-specific significance results, though only C-SIDE reports convergence status. CTSV and spVC do not include cell-type labels in their outputs, complicating the mapping of results to gene–cell-type pairs, especially in tissues with many cell types.

### S1.10. Implementation details of ctSVG methods

The implementation details of the five benchmarked ctSVG detection methods are listed as follows.

**CELINA.** We obtained *CELINA* from `https://github.com/pekjoonwu/CELINA` and ran the method by following the tutorial `https://lulushang.org/Celina_Tutorial/RCC.html`. We adopted all default parameters.

**STANCE.** We obtained *STANCE* from `https://github.com/Cui-STT-Lab/STANCE` and ran the method by following the tutorial `https://haroldsu.github.io/STANCE/tutorial.html`. We adopted all default parameters. In addition, we also followed the tutorial exactly and only passed on significant genes in stage 1 utSVG test after B–H adjustment at 0.05 FDR target to stage 2 ctSVG test.

**CTSV.** We obatined *CTSV* from `https://github.com/jingeyu/CTSV/blob/master/R/CTSV.R`. We adopted all default parameters.

**C-SIDE.** We used the `run.CSIDE.nonparam()` function from *spacexr* package acquired from `https://github.com/dmcable/spacexr` and followed the tutorial `https://raw.githack.com/dmcable/spacexr/master/vignettes/merfish_nonparametric.html`. While the recommended default number of basis functions is 15, we presented in this paper the best detection performance at 5 out of testing 5, 10 and 15. In addition, we also found increasing proportion of genes failed to converge as we increase number of bases. In addition to that, C-SIDE also implements cell type filters that only retains cell-types above a certain abundance across the tissue and within specific regions. Not only that C-SIDE fails to run in some idealized simulation settings similar as previous studies have found, but also very few cell-types can realistic tissues could pass this quality control. To ensure C-SIDE produce reasonable results, we set the cell type threshold to 0, essentially removing the cell-type level filtering mechanism.

**spVC.** We obtain *spVC* from https://github.com/shanyu-stat/spVC and followed the tutorial https://github.com/shanyu-stat/spVC/wiki/Tutorial-on-the-spVC. spVC offers two model variates: one two-step procedure that select gene-cell-type pairs that has significant constant effect and residual spatial effect to proceed to second step of testing cell-type-specific effects; a full-model version that takes just test all genes with more than 100 counts for cell-type specific effects. We choose the full model version because the two-step procedure filters out most ground truth genes in step 1 and results in nearly 0 power across all idealized simulation scenarios.

### S1.11. Adaptations in baselines of decomposed simulations

Specifically, 1) we adjusted the idealized simulation gene settings to reflect the real number of genes, ctSVGs, and an assumed 25 marker genes per cell type. To avoid the complication of maintaining equal proportion of various ctSVG signal strength, 2) we set all ctSVGs to the strongest signal strength at fold change of 4 in contrast to an equal proportion of 4, 2, 0.5, 0.25. To eliminate the confounding factor of information contained in each spot, 3) we adjusted the cell to spot ratios according to real datasets while maintaining the assumption of uniform cell density and cell-type composition. Finally, 4) we scaled tissue coordinates to match the realistic spatial range of each tissue.

### S2. Supplementary Figures

| | Random Effect Models (Fit a model for each gene-cell-type pair) | | Fixed Effect Models (Fit a model for each gene) | | |
|---|---|---|---|---|---|
| | Multiple Tests per Pair | One Test per Pair | Multiple Tests per Pair | | One Test per Pair |
| | CELINA | STANCE | CTSV | C-SIDE | spVC |
| Quanlity Control | Norm. + Relative | Norm. + utSVG test | *None* | Norm. + Relative + Cell type | Norm. |
| Constant effect | All cell types | *None* | All cell types | All cell types | All cell types |
| Spatial effect | Target cell type (11 Kernels) | All cell types (1 Kernel) | All cell types (10 Kernels) | All cell types (Penalized Spline) | All cell types (Penalized Spline) |
| P-value | Cauchy(11 Score Tests) | Score Test | Min(Cauchy(5 Wald Tests), Cauchy(5 Wald Tests)) | Min(Bon.(5 Wald Tests)) | Wald-like Test |
| Multiple Testing | *None* | *None* | Global B-H | By cell type B-H | *None* |

**Fig. S1:** Detection method summary

| | CELINA | | | STANCE | | | C−SIDE | | | CTSV | | | spVC | | |
|---|---|---|---|---|---|---|---|---|---|---|---|---|---|---|---|
| | breast | ovarian | lymph | breast | ovarian | lymph | breast | ovarian | lymph | breast | ovarian | lymph | breast | ovarian | lymph |
| Idealized | 61 | 61 | 61 | 12 | 12 | 12 | 24 | 24 | 24 | 29 | 29 | 29 | 8 | 8 | 8 |
| Baseline | 82 | 95 | 100 | 44 | 15 | 50 | 37 | 44 | 95 | 41 | 94 | 99 | 14 | 6 | 10 |
| +(i) | 66 | 73 | 79 | 43 | 20 | 100 | 31 | 23 | 97 | 37 | 100 | 100 | 4 | 7 | 10 |
| +(ii) | 92 | 99 | 50 | 52 | 32 | 27 | 41 | 55 | 94 | 49 | 54 | 100 | 40 | 29 | 8 |
| +(i,ii,iii) | 56 | 13 | 79 | 86 | 91 | 74 | 45 | 97 | 83 | 72 | 77 | 100 | 9 | 16 | 0 |
| +(iv) | 81 | 67 | 99 | 82 | 58 | 40 | 62 | 23 | 33 | 35 | 94 | 99 | 68 | 61 | 37 |
| +(i,ii,iii,iv) | 53 | 24 | 9 | 98 | 99 | 97 | 50 | 76 | 75 | 82 | 97 | 100 | 70 | 69 | 100 |
| +(ii,v) | 94 | 99 | 50 | 64 | 34 | 22 | 57 | 58 | 94 | 45 | 54 | 100 | 58 | 32 | 6 |
| +(i,ii,iii,v) | 71 | 12 | 81 | 90 | 92 | 75 | 56 | 97 | 82 | 50 | 77 | 100 | 12 | 19 | 8 |
| +(i,ii,iii,vi) | 81 | 25 | 59 | 77 | 78 | 76 | 52 | 100 | 97 | 72 | 88 | 100 | 13 | 8 | 0 |
| +All | 89 | 25 | 23 | 97 | 99 | 99 | 88 | 93 | 94 | 78 | 98 | 100 | 47 | 72 | 100 |
| Realistic | 92 | 30 | 0 | 97 | 99 | 99 | 79 | 89 | 76 | 39 | 81 | 100 | 46 | 72 | 100 |

**Fig. S2:** FDP for *decomposed simulations*.

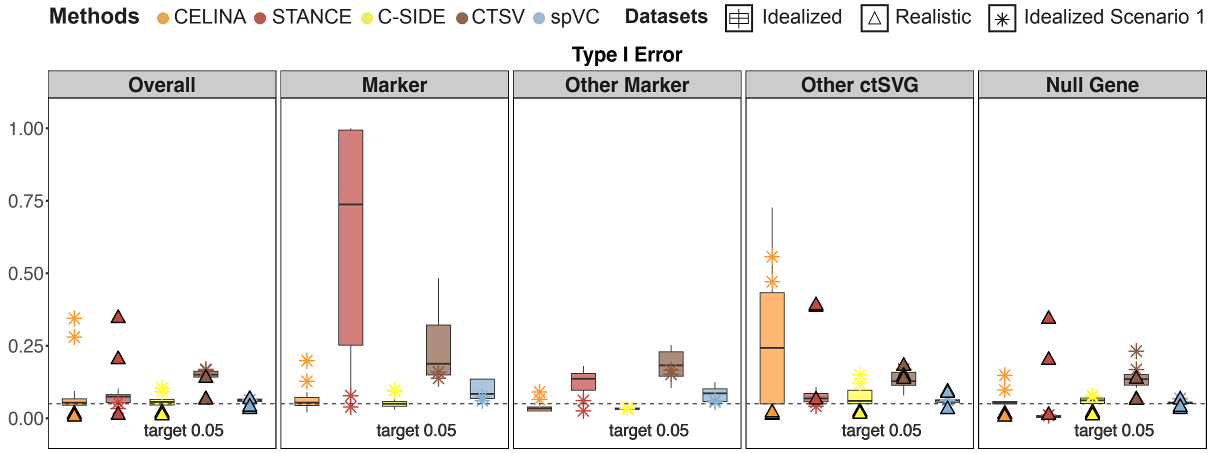


**Fig. S3:** Type I error in pure simulations stratified by the types of non-ctSVGs including tested cell type's marker genes, other cell type's marker genes, other cell type's ctSVGs and null genes.

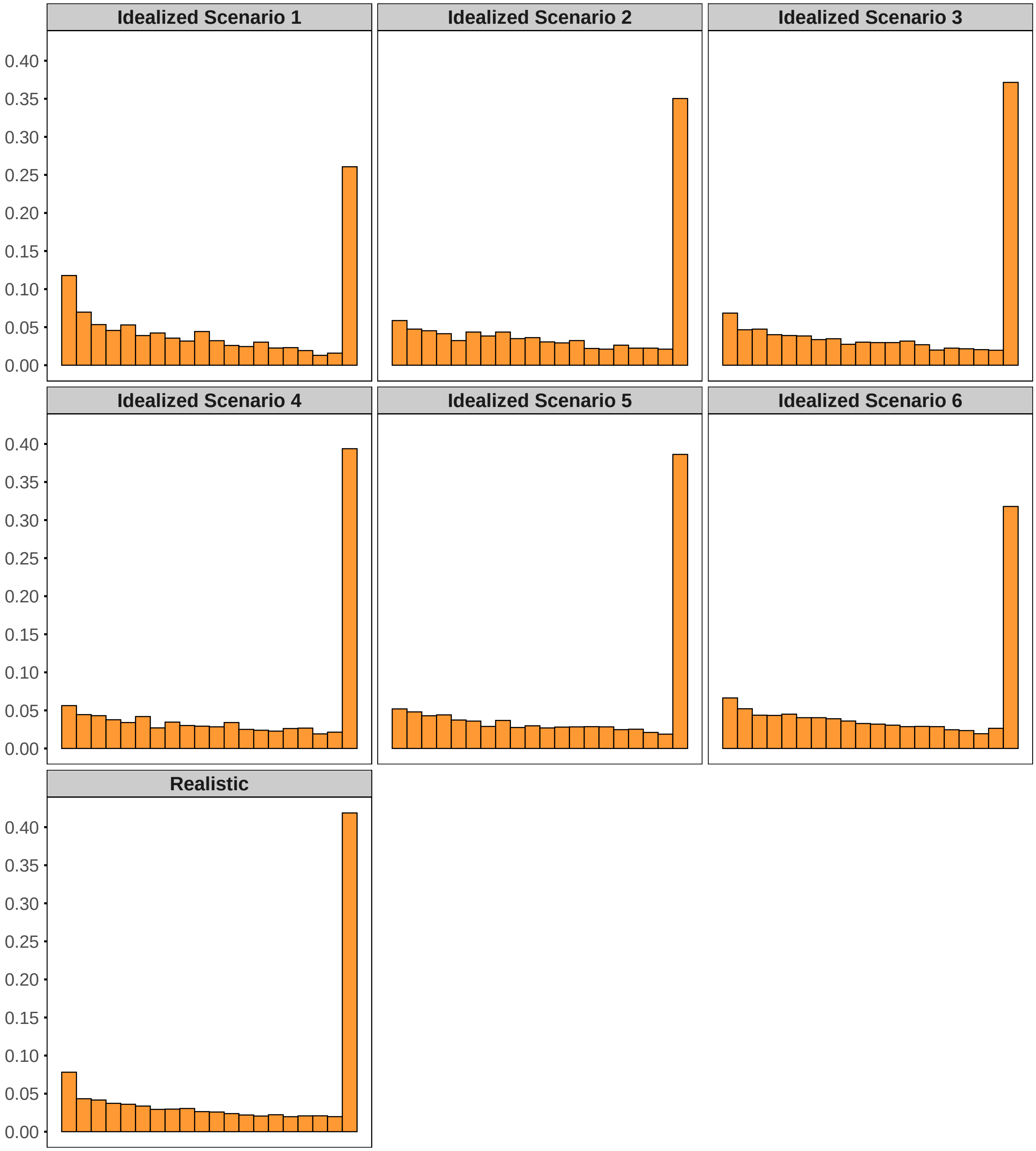


**Fig. S4:** Sample C-SIDE p value distribution for non-ctSVGs.
